# NT5DC2 downregulation suppresses monoamine oxidase activity and increases catecholamine levels in PC12D cells

**DOI:** 10.1101/2025.05.07.651779

**Authors:** Hisateru Yamaguchi, Miho Kawata, Yu Kodani, Kanako Saito, Toshiki Kameyama, Hiroshi Nagasaki, Akira Nakashima

## Abstract

**Back ground:** Genome-wide association studies have revealed the involvement of 5’-nucleotidase domain-containing protein 2 (NT5DC2) in neuropsychiatric disorders such as schizophrenia and bipolar disorder; however, its function remains unclear. Recently, we found that NT5DC2 downregulation in PC12D cells increases catecholamine synthesis by promoting the activity of tyrosine hydroxylase and that monoamine oxidase A (MAO A) might bind to NT5DC2 using Affinity Purification-Mass Spectroscopy.

**Methods and Results:** We investigated the role of NT5DC2 for MAO A activity in PC12D cells. Western blot analysis revealed that NT5DC2 primarily binds to the non-phosphorylated form of MAO A. siRNA-mediated NT5DC2 downregulation reduced MAO A activity, leading to decreased dopamine metabolism and increased noradrenaline synthesis.

**Conclusion:** Our findings suggest that NT5DC2 could affect both tyrosine hydroxylase and MAO A activity to control catecholamine synthesis. Therefore, our study provides valuable insights into disorders associated with catecholamine dysregulation, such as Parkinson’s disease and neuropsychiatric disorders.

## Introduction

According to genome-wide association studies (GWAS), 5□-nucleotidase domain-containing protein 2 (NT5DC2) is implicated in neuropsychiatric disorders such as schizophrenia, bipolar disorder, language disorder, and borderline personality disorder (Kornilov et al. 2016; Prados et al. 2015; Ripke et al. 2013; van Hulzen et al. 2017). NT5DC2 also affects the nervous system, as revealed by a Drosophila neuropsychiatric disorder-related behavioral assay (Singgih et al. 2021). These reports suggest that NT5DC2 is a key factor in the production of neurotransmitters such as dopamine, which is linked to neuropsychiatric disorders and unusual behaviors. However, although NT5DC2 has been reported to belong to the halide dehalogenase phosphatase family (Seifried et al., 2013), its function remains unknown. We used nanoscale liquid chromatography coupled to tandem mass spectrometry (nano-LC-MS/MS) to show that a rate-limiting enzyme of catecholamine, tyrosine hydroxylase (TH), binds to NT5DC2 in PC12D cells (Nakashima et al. 2019). Moreover, downregulation of NT5DC2 in PC12D cells promotes TH phosphorylation, which enhances TH activity and leads to increased catecholamine synthesis (Nakashima et al. 2020). These results indicate that NT5DC2 is a crucial key factor in catecholamine synthesis.

Recently, we overexpressed the NT5DC2-DYKDDDDK-tag (NT5DC2-tag) protein in PC12D cells and performed nano-LC-MS/MS analysis to identify proteins in the cell lysate that bind to NT5DC2 using anti-DYKDDDDK-tag affinity beads (Yamaguchi et al. 2024). The comprehensive analysis using this Affinity Purification-Mass Spectroscopy revealed that NT5DC2 has the potential to bind to 40 proteins with multiple phosphorylation sites, one of which is monoamine oxidase A (MAO A; amine oxidase [flavin-containing] A) catalyzing monoamines such as DA, noradrenaline (NAd), and adrenaline. This indicates that NT5DC2 may affect catecholamine synthesis not only through TH but also through MAO A. In this study, we examined its role of NT5DC2 for MAO A by analyzing changes in catecholamine metabolism in NT5DC2-downregulated PC12D cells.

## Materials and methods

### Cell culture

Rat pheochromocytoma PC12D cells were maintained in 24-well plates containing Dulbecco’s modified Eagle medium supplemented with 10% horse serum and 5% fetal calf serum at 37°C in humidified air containing 5% CO2, after which the cells and medium were collected (Nakashima et al. 2016). Cell lysates were obtained using a detergent or via sonication. For **Fig. 1**, the cells were suspended with phosphate-buffered saline containing 1.8 mM Ca^2+^ and 0.8 mM Mg^2+^ (PBS/Ca/Mg) and sonicated using Bioruptor UCD-250 (Diagenode Inc, Denville, NJ, USA). The cell lysates were centrifuged and the supernatants were used for experiments. For **Fig. 3** and **4**, we used Cell Lytic M (Sigma-Aldrich, St. Louis, MO, USA) containing the protease inhibitor cocktail P8340 (Sigma-Aldrich) and the phosphatase inhibitor PhosStop (Roche, Basel, Switzerland). The protein concentration was determined using a Bradford assay (Bio-Rad Laboratories, Hercules, CA, USA).

**Fig. 1.**
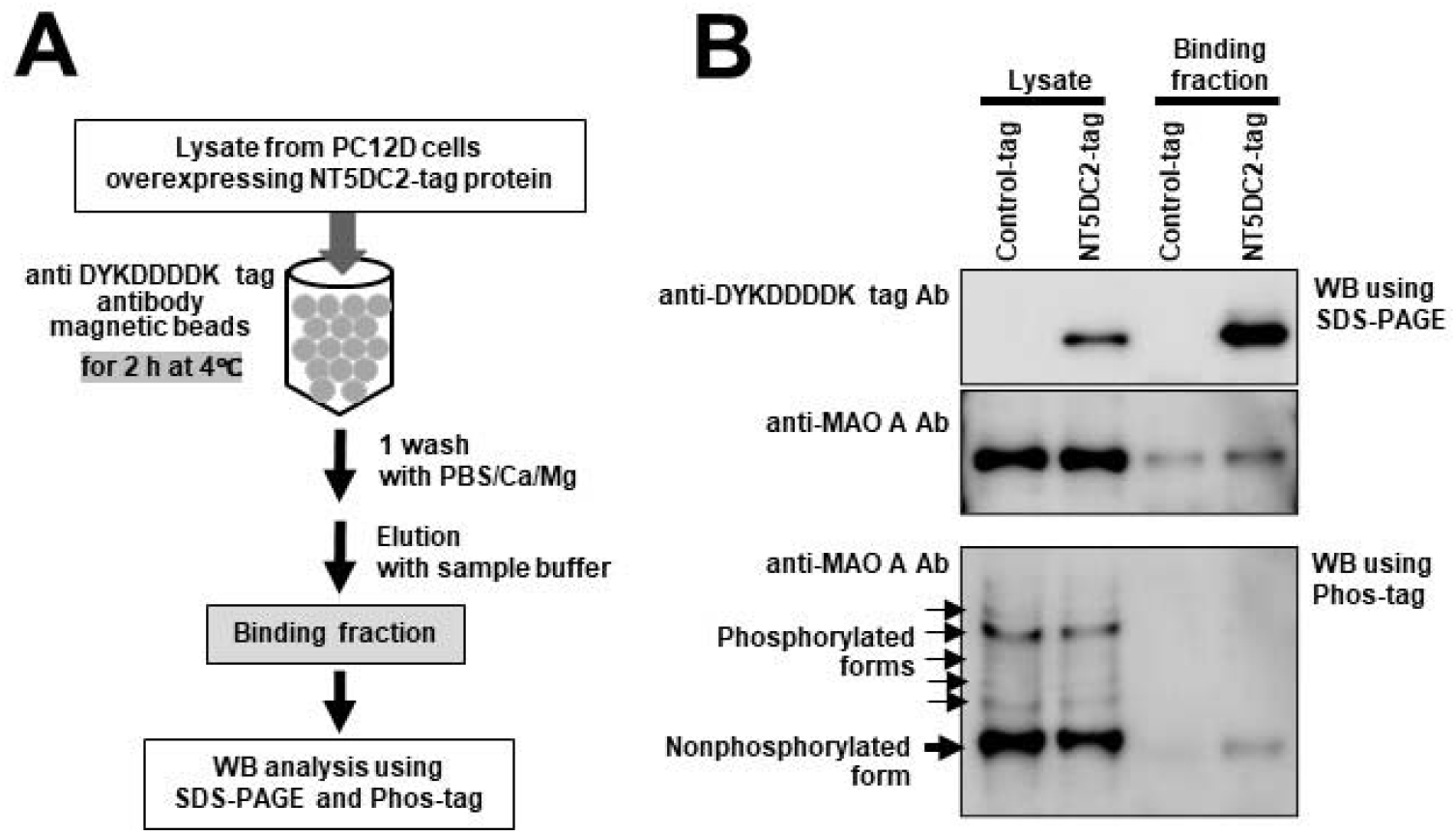
Binding of MAO A to NT5DC2. **A) Binding procedure**. Anti-DYKDDDDK-tag antibody magnetic beads were reacted with 60 µL of cell lysate (1 mg/mL protein) expressing NT5DC2-tag or control-tag at 4°C for 2 h to bind NT5DC2-tag-binding proteins. After washing with PBS/Ca/Mg, 15 µL of SDS/2-mercaptoethanol sample buffer was added, followed by incubation at 80°C for 20 min. The resulting solution was analyzed as the “binding fraction.” **B) Detection of MAO A**. Proteins in “binding fraction” were analyzed via western blotting using SDS-PAGE and Phos-tag SDS-PAGE.

### Expression vector

The pCMV6 vector, which contained rat NT5DC2 cDNA and cDNA sequences for the Myc tag and DYKDDDDK tag on its 3’ side (pCMV6-NT5DC2-DYKDDDDK-tag vector), and a control-tag vector, in which rat NT5DC2 cDNA was deleted, were constructed from the OriGene vector, as reported previously (Yamaguchi et al. 2024). Transfection was performed using Lipofectamine 3000 (Thermo Fisher Scientific, Waltham, MA, USA) following the manufacturer’s instructions. The proportions of Lipofectamine 3000 reagent, P3000 reagent, and vector were 21 µL:15 µL:7.5 µg.

### Binding of MAO A to NT5DC2

Anti-DYKDDDDK-tag antibody magnetic beads (FUJIFILM Wako, Osaka, Japan) was used to bind NT5DC2-tag protein in the cell lysate (**Fig. 1A**). The reaction was performed in a 1.5-mL PROTEOSAVE™ tube to block the nonspecific adsorption of proteins to the tube (Sumitomo Bakelite, Tokyo, Japan).

### NT5DC2 downregulation

NT5DC2 downregulation was performed using Silencer™ Select Pre-designed siRNA (Thermo Fisher Scientific, cat# s145021), with Silencer™ Select Negative Control 1 (cat# 4390843) as a negative control. PC12D cells were transfected by adding Lipofectamine RNAiMAX complex (Thermo Fisher Scientific) containing siRNA to the culture medium, as described previously (Nakashima et al. 2019) **(Fig. 2A**). The medium was refreshed after one and three days with 50 ng/mL nerve growth factor (Alomoe Labs, Jerusalem, Israel). To inhibit TH activity, 1 mM α-methyl-L-tyrosine (Watanabe Chemical, Hiroshima, Japan) was added on day three, and samples were collected on day five for DOPA analysis. For DA metabolism verification, TH inhibitor was added on day three, exogenous DA (Sigma-Aldrich) on day four, and samples were collected on day five.

**Fig. 2.**
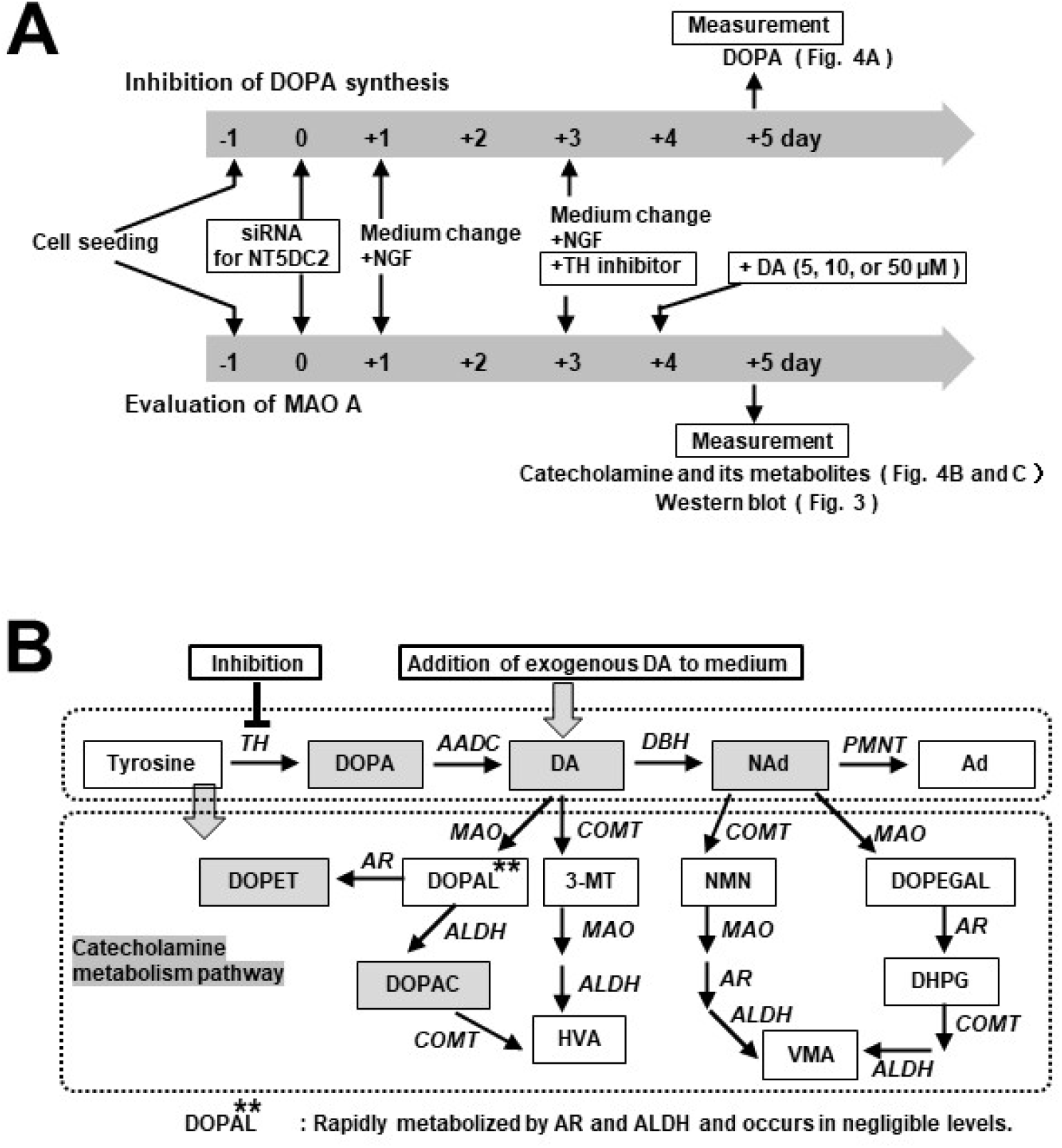
NT5DC2 downregulation on catecholamine metabolism. **A) Diagram of the experimental procedure**. The upper shows the procedure for inhibiting DOPA synthesis, while the lower shows the procedure for evaluating MAO A. **B) Catecholamine synthesis and metabolism pathway**. TH, tyrosine hydroxylase; AADC, aromatic amino acid decarboxylase; DBH, dopamine-hydroxylase; PNMT, phenylethanolamine N-methyltransferase; MAO, monoamine oxidase; AR, aldehyde reductase; ALDH, aldehyde dehydrogenase; COMT, catechol-O-methyltransferase; DOPAL, 3,4-dihydroxyphenylacetaldehyde; DOPET, 3,4-dihydroxyphenylethanol; DOPAC, 3,4-dihydroxyphenylacetic acid. Shaded squares indicate catecholamines and their metabolites identified in our experiments.

### Measurement of catecholamines and their metabolites

The lysate (3 μL) or culture medium (50 μL) of PC12D cells was added to 550 µL of 1.5 M Tris-HCl buffer (pH 8.5) containing 10 mM EDTA and 20 ng/µL α-methyldopa (Sigma-Aldrich) as an internal standard. The mixture was reacted with 10 mg of activated alumina at 4°C for 20 min, and the catecholamines and their metabolites were then released from the activated alumina through the addition of 0.2 M HCl. Catecholamines and their metabolites were measured using an HPLC HTEC-500 apparatus (EiCOM, Kyoto, Japan) equipped with an SC-5ODS column equilibrated with 80 mM citric acid-sodium acetate buffer solution (pH 2.6) containing 19% methanol, 800 mg/L 1-octanesulfonic acid sodium salt, and 5 mg/L EDTA.

### Western blot analysis

Western blotting was performed as previously described (Nakashima et al. 2016), using SuperSep™ Ace (10%) or SuperSep™ Phos-tag (7.5%) SDS-PAGE (FUJIFILM Wako). The primary antibodies used were anti-rabbit NT5DC2 (Aviva Systems Biology, San Diego, CA, USA), anti-mouse DYKDDDDK (FUJIFILM Wako), anti-rabbit MAO A (Proteintech, Rosemont, IL, USA), and anti-mouse GAPDH (FUJIFILM Wako) antibodies. The secondary antibodies used were goat anti-rabbit IgG peroxidase-conjugated IgG (Proteintech) and rabbit anti-mouse IgG peroxidase-conjugated IgG (Sigma-Aldrich). Densitometric scanning of chemiluminescence was performed on a Lumino Image Analyzer LAS-4000 (FUJIFILM, Tokyo, Japan) using Super Signal™ West Femto (Thermo Fisher Scientific).

### Statistical analysis

Statistical significance was assessed using the Student’s *t*-test or analysis of variance, followed by a post-hoc Dunnett’s *t*-test. Statistical significance was set to *p*<0.05 or *p*<0.01.

## Results

### Binding of MAO A to NT5DC2

The NT5DC2-tag protein overexpressed in PC12D cells was fixed to anti-DYKDDDDK-tag antibody magnetic beads, and then the cellular proteins bound to the NT5DC2-tag protein were analyzed (**Fig. 1A**). Western blot analysis revealed that the amount of MAO A bound to the NT5DC2-tag protein was significantly higher than that bound to the control-tag protein (control-tag, 100.0 ± 29.1%, versus NT5DC2-tag, 209.6 ± 43.0%: n=4, *p*<0.01) (**Fig. 1B**); the area of the MAO A band bound to the NT5DC2-tag protein was expressed as a percentage relative to the area of the MAO A band bound to the control-tag protein, which was set to 100%. Moreover, western blot analysis using Phos-tag SDS-PAGE revealed that approximately 50% of MAO A was phosphorylated, while NT5DC2-bound MAO A was mainly non-phosphorylated.

### Downregulation of NT5DC2

NT5DC2 downregulation was performed as shown in **Fig. 2A**, and its effect on MAO was analyzed via western blot using SDS-PAGE (**Fig. 3**). NT5DC2 downregulation did not decrease the amount of MAO A protein in PC12D cells (N = 4, *p*>0.05).

**Fig. 3.**
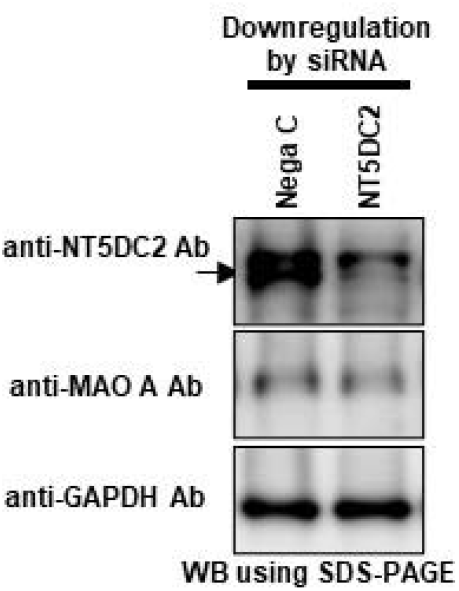
Effect of NT5DC2 downregulation on MAO A protein level. Proteins in the cell lysate were analyzed via western blotting using SDS-PAGE.

### Inhibition of DOPA synthesis

HPLC analysis primarily detected DOPA, DA, NAd, DOPET, and DOPAC, as shown in gray boxes (**Fig. 2B**). TH activity was inhibited using α-methyl-L-tyrosine to suppress DOPA synthesis in the cells (**Fig. 2A and B**). The inhibition of TH activity completely suppressed DOPA synthesis (**Fig. 4A**).

**Fig. 4.**
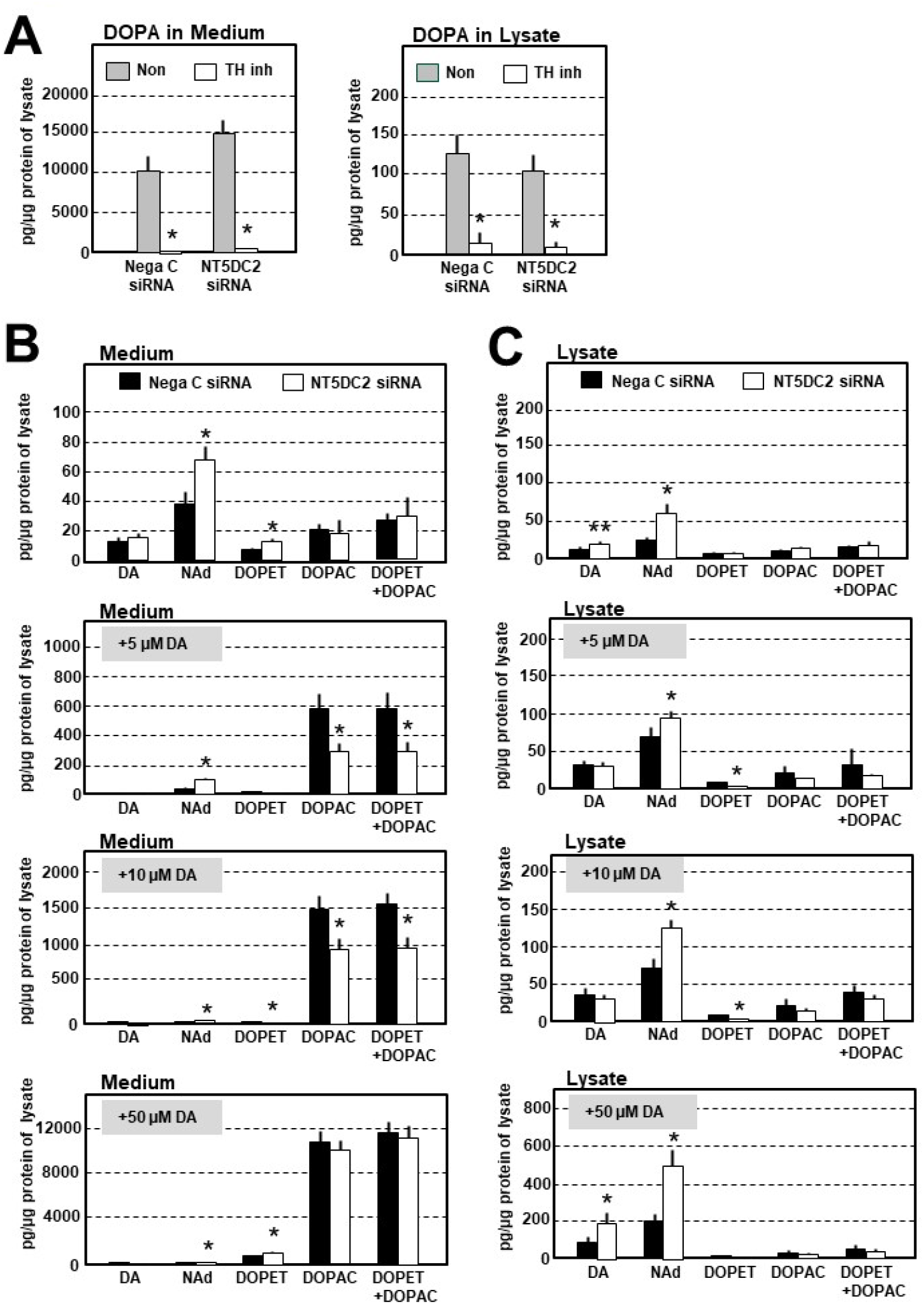
Effect of NT5DC2 downregulation on catecholamine synthesis and metabolism. **A) Inhibition of DOPA synthesis via TH inhibition**. Values of DOPA in the medium and cell lysate were corrected for the protein concentration of the cell lysate obtained from the cells in each well because they are affected by the number of cells (N = 6, *p < 0.01 vs. negative control). **B) Medium and C) Cell lysate: addition of exogenous DA to PC12D cells with inhibited DOPA synthesis**. Values of catecholamines and metabolites in the medium and cell lysate were corrected for the protein concentration of the cell lysate obtained from the cells in each well because they are affected by the number of cells (N = 8, \**p* < 0.01 or \*\**p*<0.05 vs. negative control).

### Effects of NT5DC2 downregulation on MAO activity

Among the two MAOs, MAO B normally metabolizes DA into DOPAL, whereas MAO A metabolizes DA in PC12D cells, which do not express MAO B (Binda et al. 2002; Goldstein et al. 2012). To examine NT5DC2’s effect on MAO A, we downregulated NT5DC2 in PC12D cells using siRNA, then suppressed DOPA synthesis with α-methyl-L-tyrosine and added exogenous DA to assess MAO activity (**Fig. 2A and B**). DOPAL is rapidly metabolized by ALDH and AR (Goldstein et al. 2012), allowing detection of DOPAC and DOPET but not the DA metabolite DOPAL. Therefore, we determined the sum of the amounts of DOPAC and DOPET, a value that reflects the activity of MAO A. In addition, DOPAC and DOPET are immediately released into the medium and rarely present in cells; therefore, their amounts in the medium were used for evaluation (**Fig. 4B**). In NT5DC2-downregulated PC12D cells, the amounts of DOPAC and DOPET in the medium were significantly lower than those in the control (**Fig. 4B**, when 5 and 10 μM DA were added). When 50 μM DA was added, we observed no significant difference from the control. This may be because MAO A exists in a fixed amount within the cell, resulting in saturation of the enzyme capacity. In contrast, in NT5DC2-downregulated PC12D cells, the amounts of NAd synthesized from DA in the medium and cell lysate were significantly higher than those in the control (**Fig. 4B** and **C**, when 5, 10, and 50 µM DA were added).

## Discussion

Our western blot results demonstrated that NT5DC2 binds to MAO A in PC12D cells (**Fig. 1B**), which validates the results of previous Affinity Purification-Mass Spectroscopy (Yamaguchi et al. 2024). Therefore, to estimate the role of NT5DC2 for MAO A activity, we measured DOPAC and DOPET produced by the addition of exogenous DA to PC12D cells, wherein DOPA synthesis was suppressed by the addition of a TH inhibitor. In NT5DC2-downregulated PC12D cells, the amounts of DOPAC and DOPET decreased, whereas those of NAd increased (**Fig. 4B and C**). These results indicate that NT5DC2 downregulation reduced DA metabolism by the suppression of MAO A, resulting in the promotion of NAd synthesis in cells. We previously reported that NT5DC2 downregulation promotes TH activity to increase the synthesis of DOPA and DA (Nakashima et al. 2019; Nakashima et al. 2020). Thus, our study indicates that NT5DC2 downregulation strongly promotes catecholamine synthesis in the cell by affecting MAO A in addition to TH, and therefore that NT5DC2 could control catecholamine synthesis through its impact on both TH and MAO A.

The mechanism by which NT5DC2 affects MAO A activity remains unknown. MAO A is a crucial enzyme that plays a significant role in regulating intracellular DA levels (Tábi T et al. 2020; Nagatsu and Nakashima 2022). The protein expression levels of MAO A are widely recognized as the primary determinant of its catalytic activity, as this view is supported by the majority of researchers. While there have been reports suggesting that post-translational modifications, such as phosphorylation, might contribute to its functional regulation (Mousseau and Baker 2012), only two studies have explored this possibility, presenting contradictory findings (Cao et al. 2009; Wang et al. 2009). Consequently, the nature and extent of these modifications remain largely uncertain. We have confirmed that the downregulation of NT5DC2 does not affect the intracellular levels of MAO A (**Fig. 3**). Therefore, our findings indicate that NT5DC2 plays a role in the post-translational modification of MAO A, potentially influencing its activity, and is the first identified protein involved in this process. Collectively, our study provides valuable insight into the molecular mechanisms underlying catecholamine metabolism.

## Conclusion

NT5DC2 could control catecholamine synthesis not only through its effects on TH but also via its influence on MAO A. Our study provides valuable insights into disorders associated with catecholamine dysregulation, such as Parkinson’s disease and neuropsychiatric disorders.

## Abbreviations

DA: dopamine
GWAS: Genome-wide association studies
MAO A: monoamine oxidase A
NAd: noradrenalin
NT5DC2: 5’-nucleotidase domain-containing protein 2
NT5DC2-tag: NT5DC2-DYKDDDDK-tag
TH: tyrosine hydroxylase

## Author contributions

Hisateru Yamaguchi: Investigation, Funding acquisition. Miho Kawata: Methodology. Yu Kodani: Methodology. Kanako Saito: Methodology. Toshiki Kameyama, Methodology. Hiroshi Nagasaki: Investigation. Akira Nakashima: Methodology, Investigation, Funding acquisition.

